# Xylazine’s κ opioid agonist activity is not shared with other FDA-approved α_2_-adrenergic agonists

**DOI:** 10.64898/2026.03.04.709652

**Authors:** Xi-Ping Huang, Brian E Krumm, Madigan L Bedard, Zoe A McElligott, Bryan L Roth

## Abstract

Xylazine is a α_2_-adrenergic agonist typically used in as a sedative and analgesic in veterinary medicine. For some years, xylazine has been reported as an additive to fentanyl on the illicit drug market and has been associated with severe side-effects including severe ulcerations and potential amputations at the sites of injection along with an increased risk of respiratory depression and death. We recently reported that xylazine has modest κ opioid agonist activity *in vitro* and *in vivo* and asked if other α_2_-adrenergic agonists had similar off-target activities. To test this hypothesis, we profiled US FDA-approved α_2_-adrenergic agonists at 320 G protein coupled receptors (GPCRs) to identify potentially deleterious and/or beneficial off-targets. Although all other tested α_2_-adrenergic agonists were devoid of κ opioid agonist activity, each had a distinct pattern of activity at various GPCRs and differential patterns of signaling bias at α_2-_receptor subtypes. These findings suggest potential molecular targets for both side-effects and therapeutic activities among known α_2_-adrenergic agonists.

## Introduction

Xylazine is an α_2_-adrenergic agonist approved in veterinary medicine for sedation and analgesia^1,2^. Some years ago, the combinations of xylazine and fentanyl and xylazine and heroin were reported to be associated with increased risk of overdose^3^. Later, initial reports appeared associating xylazine/fentanyl combinations with ulcerations and necrotic tissue at the sites of injection^4^, while recent studies have found that xylazine-associated wounds also are present distal to injection sites ^5^. This phenomenon of ulcerations and necrosis associated with xylazine injections and other routes of administration ^6 6 5^is now recognized as a *bona fide* clinical syndrome ^7^. In addition to the risk of localized tissue necrosis, there are also concerns that xylazine may potentiate the deleterious actions of fentanyl ^8,9^. Because of the emerging public health crises of this unregulated drug mixture, the FDA has issued an alert to physicians and other health-care providers regarding the potential dangers of xylazine and xylazine/fentanyl combinations (https://www.fda.gov/drugs/drug-safety-and-availability/fda-alerts-health-care-professionals-risks-patients-exposed-xylazine-illicit-drugs).

It has been assumed that the actions of xylazine are due solely to its α_2_-adrenergic agonist activity, although we recently reported that xylazine is also a low potency κ opioid receptor (KOR) agonist ^10^ in mice *in vitro* and *in vivo*. These findings have raised the intriguing question as to whether other α_2_-adrenergic agonists have activity at KOR and/or other molecular targets. To address this possibility, we have profiled several FDA approved α_2_-adrenergic agonists at 320 GPCRs using our PRESTO-Tango platform ^11^. We find that although xylazine is unique in its weak KOR agonist activity, every tested α_2_-adrenergic agonist had significant off-target actions at other GPCRs, some of which could be clinically relevant. Additionally, we find that the patterns of signaling bias downstream of various α_2_-adrenergic subtypes is highly ligand-dependent. We find xylazine, for instance, is highly G protein biased compared with other tested α_2_-agonists.

## Results

### Profiling α_2_-AR agonists against 320 GPCRs reveals potential off-targets

In initial studies, we screened 10 FDA-approved α_2_-adrenergic agonists (approved for human use: apraclonidine, clonidine, dexmedetomidine, guanfacine, lofexidine, medetomidine, and tizanidine; approved for veterinary use: detomidine, romifidine, and xylazine) and the mixed opioid agonist pentazocine (which is also associated with skin lesions ^12,13^) at 320 druggable GPCRs using our PRESTO-Tango resource ^11^ at 10 μM final concentration. We also examined two known and commercially available metabolites of xylazine: 3-OH-xylazine and 4-OH-xylazine ^14^. As anticipated (Fig 1) we found all 9 tested α_2_-adrenergic agonists along with 3-OH-xylazine and 4-OH-xylazine activated α_2A_-, α_2B_-, and α_2C_ -adrenergic receptors. Pentazocine had no agonist activity at α_2_-adrenergic receptors while activation of μ and κ-opioid (KOR) receptors was measured (Fig 1). Specifically, Xylazine activated KOR and D_2_ as reported previously ^10^. Apraclonidine activated D_2_, D_3_, and 5-HT_1F_. Detomidine activated D_2_, Dexmedetomidine activated D_2_ and 5-HT_5A_, Guanfacine activated D_2_, 5-HT_1A_, 5-HT_1D_, 5-HT_2A_, and 5-HT_2B._ and Lofexidine activated D_2_, 5-HT_1A_ and 5-HT_1F_.

**Fig 1.**
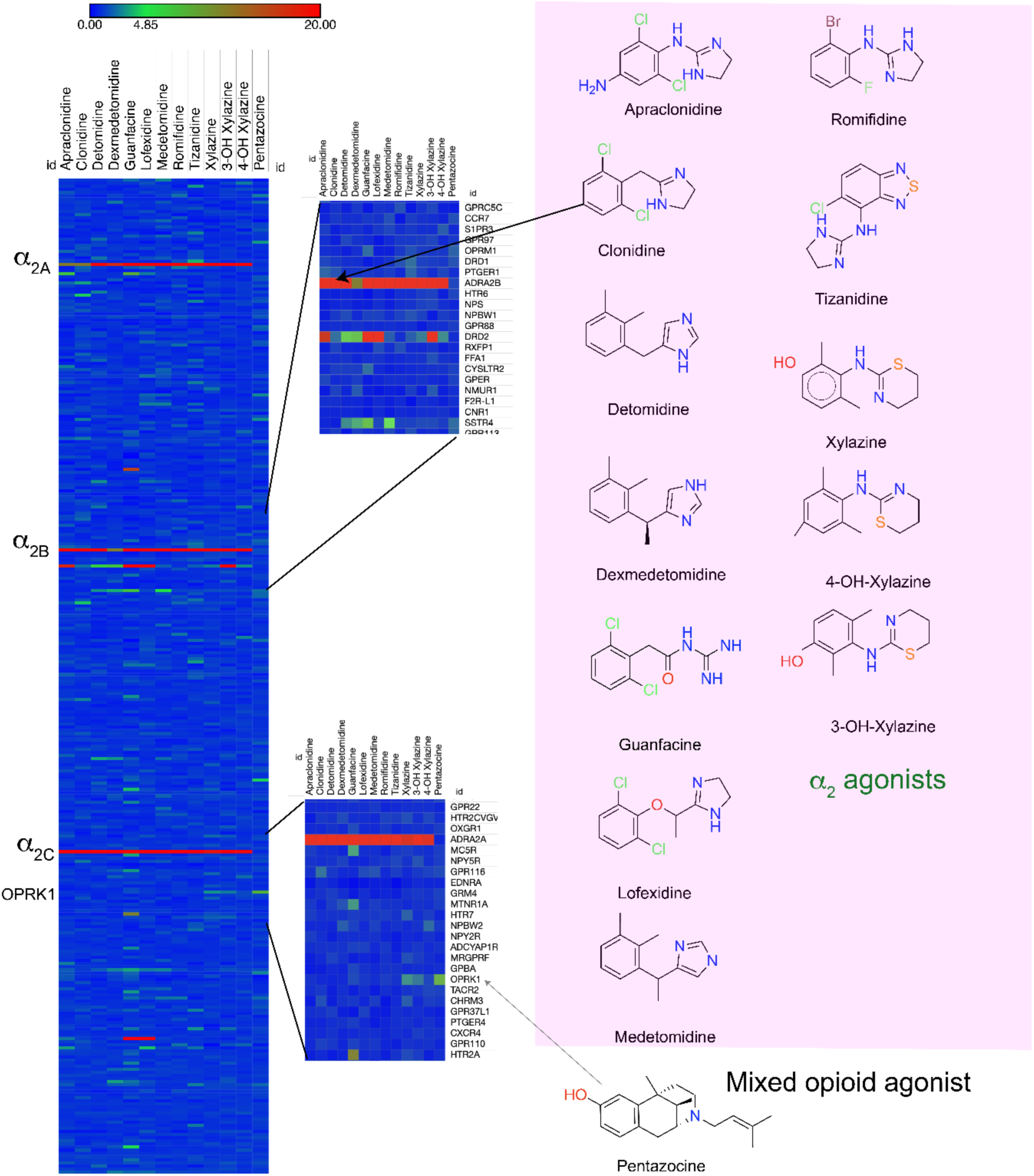
Heatmap presentation of agonist activity of FDA approved Alpha2A agonists at human GPCRome. Activity represented mean fold of basal from 4 wells with sample (10 μM) over average basal of 4 wells with medium alone. Xylazine metabolites, 3-OH Xylazine and 4-OH Xylazine, and mixed opioid agonist (-)-Pentazocine were also included for comparison. Chemical structures were shown on the right.

As shown in Supplementary Figure 1 and Supplementary Table 1, several α_2A_-AR agonists were confirmed to have agonist activity at other GPCRs, including D_2_- and D_3_-dopamine and 5-HT_1A_- and 5-HT_1F_-serotonin receptors. We next performed additional functional assays to examine and compare their agonist activity for inhibition of cAMP production and GoA dissociation using BRET assays, as these four receptors are Gi/o-coupled GPCRs. As shown in Figure 2 and Table 1, Lofexidine was confirmed to have full agonist activity with comparable potency values in both assays at 5-HT_1A_ - serotonin receptors. Apraclonidine and Lofexidine were full agonists with similar potency values from 2 functional readouts at 5-HT_1F_ -serotonin receptors. At D_2_-dopamine receptors, Apraclonidine, Detomidine, and Lofexidine were low potency, weak partial agonists, while at D_3_-dopamine receptors, Apraclonidine was a low potency, weak partial agonist.

**Fig 2.**
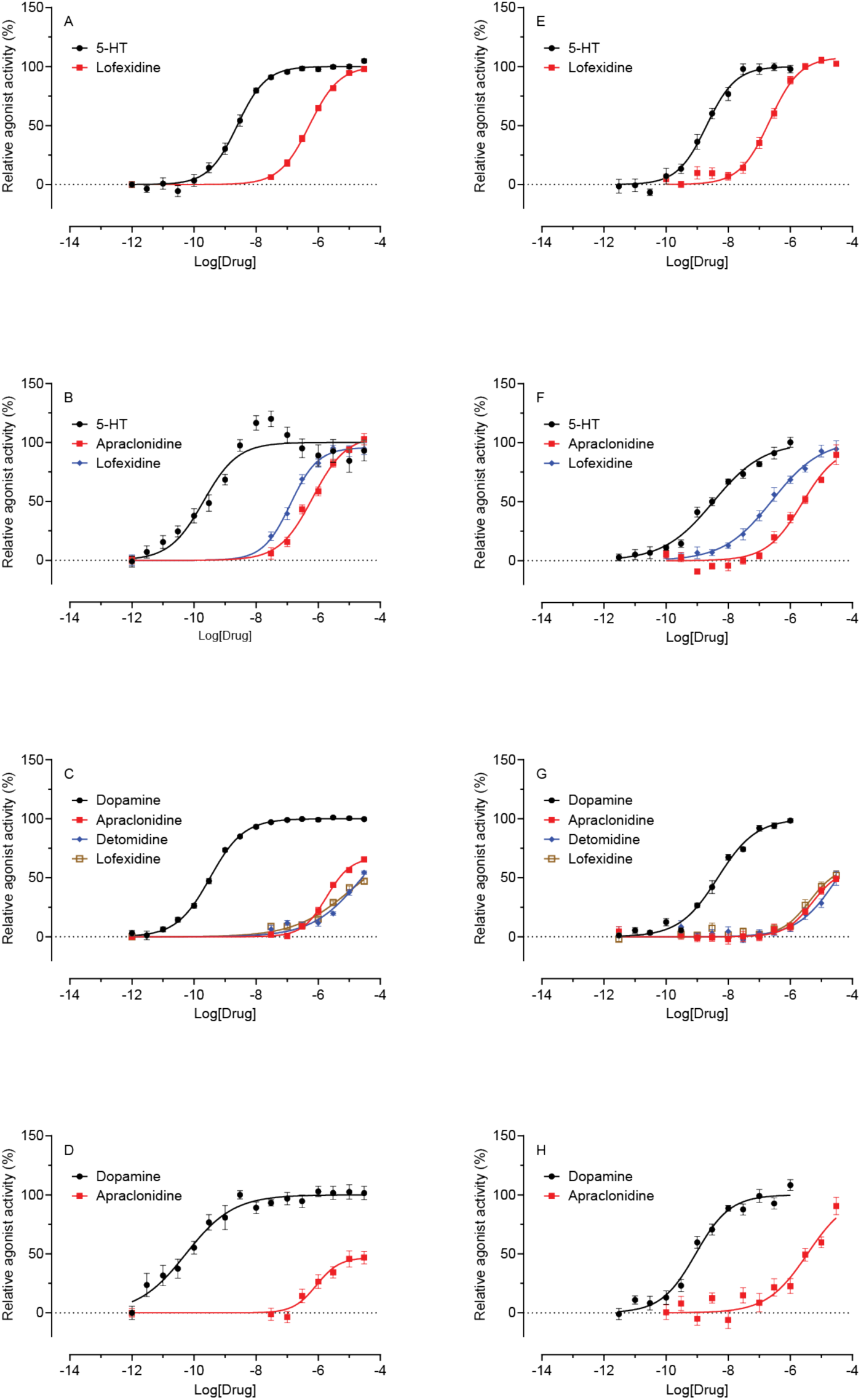
Agonist activity of selected compounds at selected GPCRs. Agonist activity was tested in both GloSensor cAMP assays (A, B, C, and D) and GoA dissociation BRET assays (E, F, G, and H) in transiently transfected HEK293 T cells. Results represented means ± SEM from a minimum of 3 independent assays, each in quadruplicate, and normalized to corresponding reference agonists (5-HT for 5-HT receptors and Dopamine for Dopamine receptors) and pooled for nonlinear fitting to the built-in four-parameter logistic function in Prism (V10). Pharmacological parameters were extracted and summarized in Table 1.

**Table 1.** Pharmacological parameters of selected compounds at selected GPCRs in the inhibition of cAMP production and GoA dissociation assays. Values were extracted from fitting normalized results with the built-in four-parameter logistic function in the Prism and represented mean ± SEM from a minimum of 3 independent assays, each in quadruplicate set. Concentration response curves are presented in Figure 5. Note, * indicates that E_max_ was constrained to be 100% for stable fitting.

| Compound<sup>19</sup> | E<sub>max</sub> (%) | pEC<sub>50</sub> | Hill slope |
|---|---|---|---|
| Inhibition of cAMP production (GloSensor cAMP assays) |  |  |  |
| 5-HT at 5-HT<sub>1A</sub> | 100.1 ± 1.4 | 8.61 ± 0.04 | 0.94 ± 0.08 |
| Lofexidine at 5-HT<sub>1A</sub> | 100.4 ± 2.42 | 6.28 ± 0.04 | 0.90 ± 0.06 |
| 5-HT at 5-HT<sub>1F</sub> | 100.0 ± 2.8 | 9.70 ± 0.10 | 0.79 ± 0.13 |
| Apraclonidine at 5-HT<sub>1F</sub> | 105.9 ± 5.0 | 6.18 ± 0.09 | 0.81 ± 0.10 |
| Lofexidine at 5-HT<sub>1F</sub> | 95.7 ± 2.9 | 6.89 ± 0.06 | 0.98 ± 0.14 |
| Dopamine at D<sub>2</sub> | 100.0 ± 0.7 | 9.48 ± 0.02 | 0.80 ± 0.03 |
| Apraclonidine at D<sub>2</sub> | 67.4 ± 3.1 | 5.73 ± 0.06 | 1.11 ± 0.13 |
| Detomidine at D<sub>2</sub> | 100.0 ± 44.7 | 4.59 ± 0.70 | 0.54 ± 0.12 |
| Lofexidine at D<sub>2</sub> | 76.6 ± 16.6 | 5.01 ± 0.42 | 0.49 ± 0.07 |
| Dopamine at D<sub>3</sub> | 100.0 ± 2.5 | 10.26 ± 0.11 | 0.57 ± 0.08 |
| Apraclonidine at D<sub>3</sub> | 46.9 ± 5.1 | 6.06 ± 0.17 | 1.17 ± 0.43 |
| GoA dissociation (BRET assays) |  |  |  |
| 5-HT at 5-HT<sub>1A</sub> | 100.0 ± 3.1 | 8.70 ± 0.07 | 0.93 ± 0.12 |
| Lofexidine at 5-HT<sub>1A</sub> | 108.0 ± 3.5 | 6.66 ± 0.06 | 0.89 ± 0.09 |
| 5-HT at 5-HT<sub>1F</sub> | 100.0 ± 4.2 | 8.48 ± 0.11 | 0.54 ± 0.05 |
| Apraclonidine at 5-HT<sub>1F</sub> | 97.5 ± 10.6 | 5.62 ± 0.17 | 0.76 ± 0.12 |
| Lofexidine at 5-HT<sub>1F</sub> | 102.2 ± 6.2 | 6.58 ± 0.14 | 0.55 ± 0.06 |
| Dopamine at D<sub>2</sub> | 100.0 ± 2.5 | 8.34 ± 0.06 | 0.70 ± 0.05 |
| Apraclonidine at D<sub>2</sub> | 57.8 ± 11.9 | 5.30 ± 0.24 | 0.99 ± 0.28 |
| Detomidine at D<sub>2</sub> | 100.0* | 4.48 ± 1.13 | 0.67 ± 0.26 |
| Lofexidine at D<sub>2</sub> | 60.8 ± 10.4 | 5.37 ± 0.21 | 0.94 ± 0.23 |
| Dopamine at D<sub>3</sub> | 100.0 ± 3.0 | 9.05 ± 0.08 | 0.78 ± 0.10 |
| Apraclonidine at D<sub>3</sub> | 100.0* | 5.45 ± 0.40 | 0.66 ± 0.19 |

### Xylazine is the unique among α_2_-AR agonists as a κ−opioid receptor agonist

Xylazine was confirmed to be a KOR agonist as we previously reported ^10^. Here we tested 9 other α_2_-AR agonists at KOR for potential agonist activity. First, to directly measure KOR mediated inhibitory G-protein activity, we compared their activity in inhibiting cAMP production in the transiently transfected HEK293 T cells using the GloSensor cAMP assays. As shown in Figure 3 and Table 2, and consistent with previously reported agonist activity, xylazine showed G_i/o_ agonist activity at KOR but not at control HEK293 T cells. Apraclonidine, detomidine, and medetomidine displayed weak activity at HEK293 T cells with or without KOR transfections, indicating nonspecific activities at high concentrations. We then carried out KOR mediated G_oA_ dissociation activity in BRET assays (Fig 3C and Table 2) and xylazine was confirmed to be a KOR agonist, while other α_2_-AR agonists showed minimal (Lofexidine and Detomidine) or no agonist activity.

**Fig 3.**
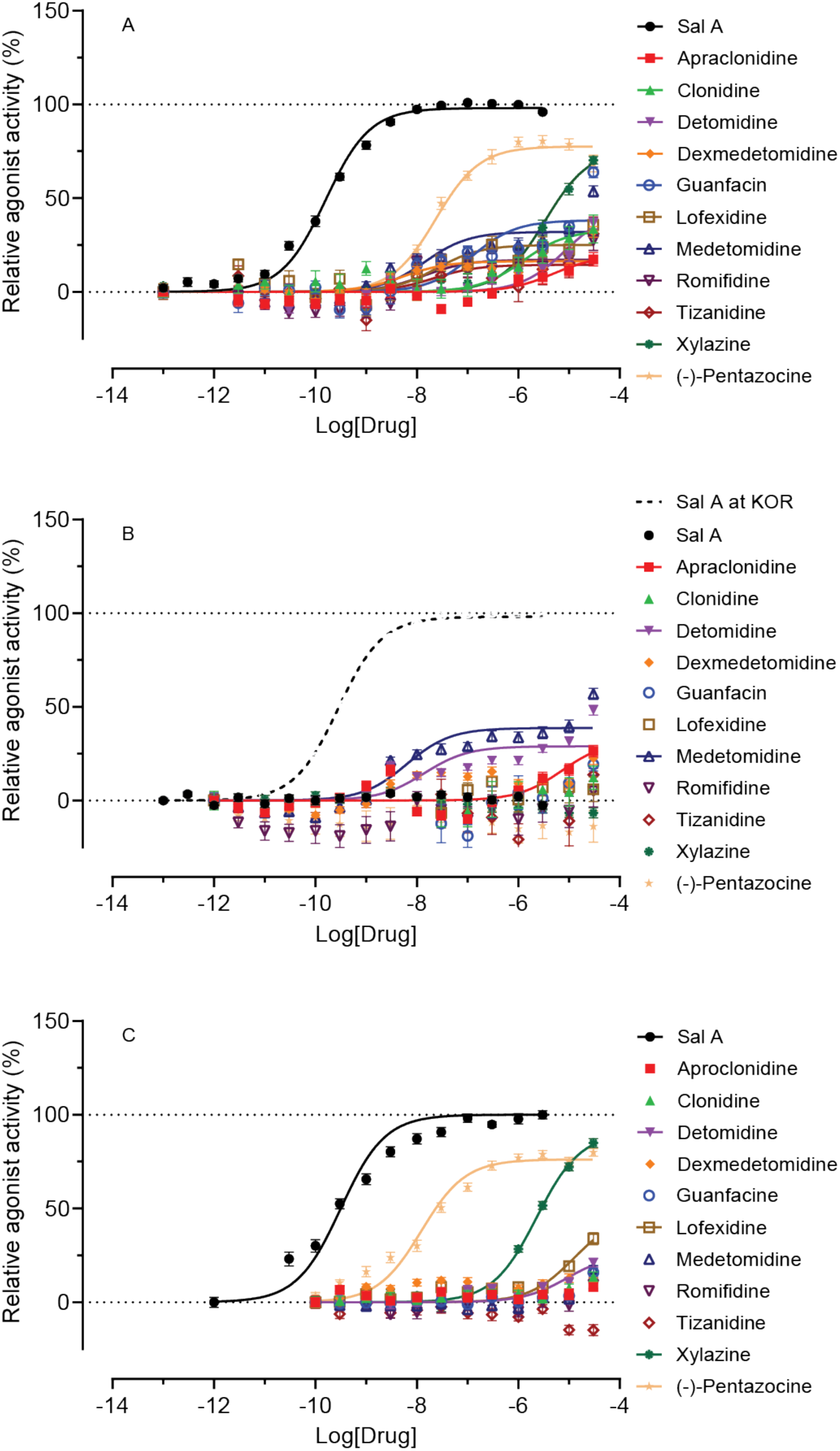
Agonist activity of FDA approved alpha2A agonists at KOR receptors. GloSensor cAMP activity in HEK293 T cells transfected with KOR (A) or without KOR (B), and KOR mediated GoA dissociation assays (BRET2) (C). Results were normalized to KOR agonist Sal A, represented means ± SEM from a minimum of 3 independent assays, each in quadruplicate set, and pooled for fitting to the built-in three-parameter logistic function in the Prism (V10). Pharmacological parameters were extracted and summarized in Table 21. For activity at control HEK293 T cells (Panel B), results were first normalized to the fold of basal and re-normalized to the Sal A activity at KOR in percentage for comparison.

**Table 2.** Gi GloSensor cAMP summary at KOR. Parameters were extracted from fitting pooled normalized results to the built-in three-parameter logistic function in the Prism and represented mean ± SEM from a minimum of 3 assays, each in quadruplicate set, as in Figure 3A and 32C. Note: NA for not available due to unstable fitting.

| **Compounds** | **G<sub>i</sub> GloSensor cAMP** |  | **G<sub>oA</sub> dissociation** |  |
|---|---|---|---|---|
|  | **E<sub>max</sub>** | **pEC<sub>50</sub>** | **E<sub>max</sub>** | **pEC<sub>50</sub>** |
| Sal A | 98.1 ± 0.7 | 9.80 ± 0.02 | 98.6 ± 1.5 | 9.53 ± 0.05 |
| Apraclonidine | 19.3 ± 4.7 | 5.31 ± 0.32 | NA | NA |
| Clonidine | 33.0 ± 4.6 | 5.91 ± 0.25 | NA | NA |
| Detomidine | 44.0 ± 5.2 | 5.08 ± 0.14 | 25.9 ± 6.3 | 5.04 ± 0.28 |
| Dexmededomidine | 16.3 ± 0.9 | 8.28 ± 0.16 | NA | NA |
| Guanfacine | 38.2 ± 2.6 | 6.91 ± 0.17 | NA | NA |
| Lofexidine | 25.0 ± 2.3 | 7.61 ± 0.25 | 46.7 ± 8.6 | 4.91 ± 0.19 |
| Medetomidine | 32.1 ± 1.5 | 7.84 ± 0.13 | NA | NA |
| Romifidine | 17.2 ± 1.8 | 7.87 ± 0.29 | NA | NA |
| Tizanidine | 14.4 ± 2.7 | 7.68 ± 0.52 | NA | NA |
| Xylazine | 76.2 ± 4.1 | 5.47 ± 0.08 | 90.5 ± 2.1 | 5.64 ± 0.04 |
| (-)-Pentazocine | 77.6 ± 1.5 | 7.63 ± 0.05 | 76.1 ± 1.1 | 7.91 ± 0.04 |

### Comparative α_2_-AR agonist activity

We then compared agonist activities of all 9 α_2_-AR agonists and 2 xylazine metabolites, 3-OH xylazine and 4-OH xylazine, at the 3 α_2_-AR subtypes (α_2A_-AR, α_2B_-AR, α_2C_-AR) for their inhibitory G protein activity by measuring cAMP production in transiently transfected HEK293 T cells (Figure 4A, 4B, 4C and Table 3). The endogenous adrenergic agonist norepinephrine was included as a reference agonist. In general, these drugs all showed potent and efficacious agonist activity, with an efficacy trend lower at α2C subtype and with a potency trend higher at α_2A_-AR than at α_2B_-AR and/or α_2C_-AR. Detomidine and medetomidine, however, were more potent at α2B than at α2A or α2C. Lofexidine had similar potency at all three subtypes. Dexmedetomidine is the most potent agonist, while xylazine is the least potent agonist, at each subtype among all tested compounds.

**Fig 4.**
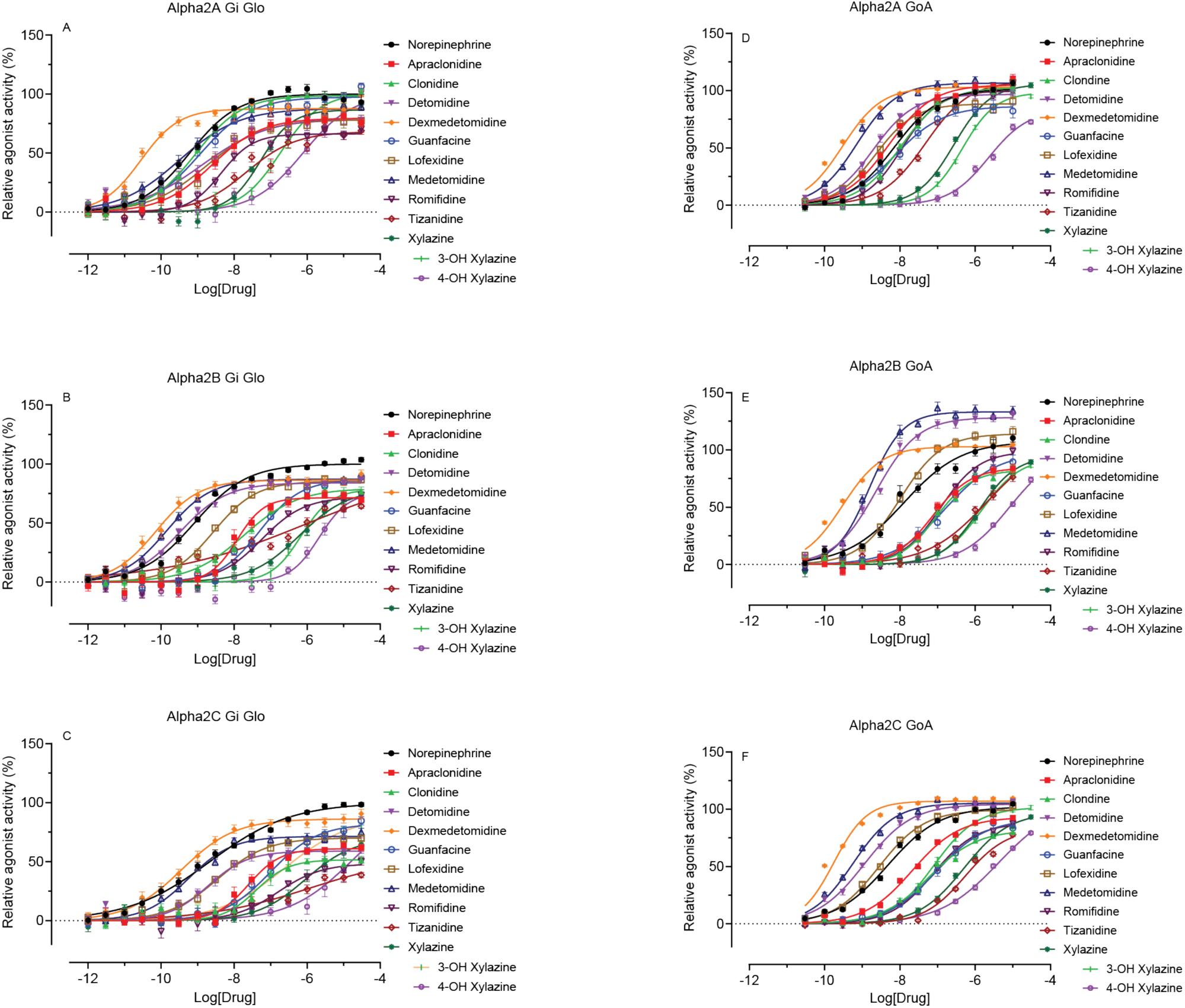
Comparison of agonist activity of FDA approved Alpha2A agonists at Alpha2A (A, D), Alpha2B (B, E), and Alpha2C (C, F) in GloSensor cAMP assays (A, B, C) and GoA dissociation BRET2 assays (D, E, F). The assays were performed in transiently transfected HEK293 T cells. Results were normalized to reference agonist Norepinephrine, represented means ± SEM from a minimum of 3 independent assays, each in quadruplicate set, and pooled for nonlinear fitting to the built-in four-parameter logistic function in Prism (V10). Pharmacological parameters were extracted from nonlinear fitting and summarized in Table 3 and 4.

**Table 3.** Pharmacological parameters of Alpha2 mediated inhibition of cAMP production measured in GloSensor cAMP assays. Values were extracted from fitting normalized results with the built-in four-parameter logistic function in the Prism and represented mean ± SEM from a minimum of 3 independent assays, each in quadruplicate set. Concentration response curves are presented in Figure 4A, 4B, and 4C. Note: * Emax was constrained to be less than 100% for stable fitting.

| Compounds | E<sub>max</sub> (%) | pEC<sub>50</sub> | Hill | ΔLog(E<sub>max</sub>/EC<sub>50</sub>) |
|---|---|---|---|---|
| Alpha2A |  |  |  |  |
| Norepinephrine | 100.0 ± 1.7 | 9.22 ± 0.06 | 0.66 ± 0.05 | 0.00 |
| Apraclonidine | 79.5 ± 2.0 | 8.61 ± 0.09 | 0.63 ± 0.07 | 0.51 |
| Clonidine | 98.7 ± 1.5 | 9.05 ± 0.06 | 0.68 ± 0.05 | 0.16 |
| Detomidine | 79.7 ± 2.9 | 8.88 ± 0.14 | 0.48 ± 0.06 | 0.24 |
| Dexmedetomidine | 87.5 ± 1.4 | 10.56 ± 0.07 | 0.82 ± 0.09 | -1.40 |
| Guanfacine | 97.4 ± 2.5 | 8.94 ± 0.09 | 0.62 ± 0.07 | 0.24 |
| Lofexidine | 78.2 ± 2.3 | 8.76 ± 0.11 | 0.56 ± 0.06 | 0.35 |
| Medetomidine | 87.7 ± 1.8 | 9.50 ± 0.08 | 0.53 ± 0.05 | -0.34 |
| Romifidine | 66.2 ± 1.7 | 8.35 ± 0.08 | 1.00 ± 0.15 | 0.69 |
| Tizanidine | 68.1 ± 2.7 | 7.65 ± 0.12 | 0.65 ± 0.09 | 1.40 |
| Xylazine | 86.5 ± 2.5 | 7.28 ± 0.07 | 1.01 ± 0.13 | 1.88 |
| 3-OH Xylazine | 100.6 ± 4.2 | 6.73 ± 0.09 | 0.79 ± 0.10 | 2.49 |
| 4-OH Xylazine | 100.0 ± 9.4 | 6.11 ± 0.19 | 0.64 ±0.11 | 3.11 |
| Alpha2B |  |  |  |  |
| Norepinephrine | 100.0 ± 1.3 | 9.09 ± 0.05 | 0.60 ± 0.03 | 0.00 |
| Apraclonidine | 71.4 ± 2.0 | 7.92 ± 0.07 | 1.18 ± 0.19 | 1.02 |
| Clonidine | 79.4 ± 2.0 | 7.73 ± 0.08 | 0.59 ± 0.05 | 1.26 |
| Detomidine | 84.0 ± 1.1 | 9.49 ± 0.05 | 0.67 ± 0.05 | -0.28 |
| Dexmedetomidine | 86.3 ± 1.4 | 10.14 ± 0.06 | 0.74 ± 0.07 | -1.11 |
| Guanfacine | 86.8 ± 1.9 | 7.29 ± 0.06 | 0.77 ± 0.06 | 1.74 |
| Lofexidine | 85.4 ± 1.5 | 8.56 ± 0.06 | 0.74 ± 0.06 | 0.46 |
| Medetomidine | 87.1 ± 0.9 | 9.84 ± 0.04 | 0.72 ± 0.04 | -0.81 |
| Romifidine | 72.1 ± 2.5 | 7.25 ± 0.09 | 0.74 ± 0.09 | 1.70 |
| Tizanidine | 97.7 ± 20.4 | 6.05 ± 0.89 | 0.22 ± 0.03 | 3.03 |
| Xylazine | 78.7 ± 5.4 | 6.21 ± 0.14 | 0.65 ± 0.09 | 2.78 |
| 3-OH Xylazine | 78.4 ± 3.8 | 6.16 ± 0.08 | 1.06 ± 0.16 | 2.82 |
| 4-OH Xylazine | 75.8 ± 6.4 | 5.67 ± 0.11 | 1.09 ± 0.23 | 3.30 |
| Alpha2C |  |  |  |  |
| Norepinephrine | 100.0 ± 2.1 | 8.67 ± 0.08 | 0.39 ± 0.02 | 0.00 |
| Apraclonidine | 61.1 ± 1.8 | 7.61 ± 0.08 | 0.98 ± 0.14 | 0.85 |
| Clonidine | 51.6 ± 1.7 | 7.40 ± 0.08 | 0.98 ± 0.14 | 0.98 |
| Detomidine | 58.9 ± 1.6 | 8.71 ± 0.09 | 0.80 ± 0.11 | -0.27 |
| Dexmedetomidine | 86.2 ± 1.7 | 9.44 ± 0.08 | 0.59 ± 0.05 | -0.83 |
| Guanfacine | 82.0 ± 3.2 | 7.01 ± 0.10 | 0.67 ± 0.08 | 1.57 |
| Lofexidine | 70.2 ± 1.7 | 8.46 ± 0.08 | 0.64 ± 0.06 | 0.06 |
| Medetomidine | 71.4 ± 1.2 | 9.25 ± 0.06 | 0.73 ± 0.06 | -0.73 |
| Romifidine | 48.4 ± 3.3 | 6.80 ± 0.16 | 0.73 ± 0.15 | 1.55 |
| Tizanidine | 55.2 ± 16.5 | 5.93 ± 0.96 | 0.31 ± 0.09 | 2.48 |
| Xylazine | 70.2 ± 6.5 | 6.04 ± 0.18 | 0.65 ± 0.11 | 2.48 |
| 3-OH Xylazine | 88.0 ± 7.3 | 6.64 ± 0.22 | 0.48 ± 0.07 | 1.97 |
| 4-OH Xylazine | 100.0* | 4.75 ± 1.40 | 0.40 ±0.13 | 3.92 |

As shown in Figure 3B, in our cAMP assays we found some compounds may be affected by nonspecific activity at control cells in the absence of transfected receptors. We thus compared their agonist activity in receptor specific G_oA_ dissociation BRET assays (Figure 43D, 4E, and 4F and Table 4) with norepinephrine served as reference agonist. Here we focused on G_oA_ as it is the main inhibitory G protein expressed in the brain. As in the cAMP assays, all tested α2 agonists demonstrated strong agonist activity in the G_oA_ dissociation assay but without the same efficacy and or potency trends from the cAMP assays. Relative efficacy values were greater than 83% with apraclonidine and clonidine being more potent at α2A than at α2B or α2C. Detomidine and medetomidine had similar potency values at all three subtypes. As in the cAMP assays, dexmedetomidine was the most potent agonist, while xylazine was the least potent agonist among all tested compounds.

**Table 4.** Pharmacological parameters of Alpha2 mediated GoA dissociation measured in the TRUPATH assays. Values were extracted from fitting normalized results with the built-in four-parameter logistic function in the Prism and represented mean ± SEM from a minimum of 3 independent assays, each in quadruplicate set. Concentration response curves are presented in Figure 4D, 4E, and 4F.

| **Compounds** | **E<sub>max</sub> (%)** | **pEC<sub>50</sub>** | **Hill** | **ΔLog(E<sub>max</sub>/EC<sub>50</sub>)** |
|---|---|---|---|---|
| Alpha2A |  |  |  |  |
| Norepinephrine | 101.3 ± 1.5 | 8.15 ± 0.04 | 0.73 ± 0.04 | 0.00 |
| Apraclonidine | 105.1 ± 1.7 | 8.30 ± 0.05 | 0.74 ± 0.05 | -0.13 |
| Clonidine | 102.5 ± 1.8 | 7.97 ± 0.05 | 0.77 ± 0.05 | 0.19 |
| Detomidine | 96.7 ± 1.5 | 8.71 ± 0.05 | 0.79 ± 0.06 | -0.58 |
| Dexmedetomidine | 102.9 ± 0.6 | 9.52 ± 0.02 | 0.83 ± 0.03 | -1.36 |
| Guanfacine | 86.4 ± 3.1 | 8.14 ± 0.10 | 0.68 ± 0.09 | -0.06 |
| Lofexidine | 88.5 ± 1.4 | 8.65 ± 0.05 | 0.83 ± 0.07 | -0.56 |
| Medetomidine | 106.6 ± 1.2 | 9.15 ± 0.04 | 0.82 ± 0.05 | -0.98 |
| Romifidine | 99.6 ± 1.5 | 7.94 ± 0.04 | 0.89 ± 0.06 | 0.20 |
| Tizanidine | 108.3 ± 2.2 | 7.32 ± 0.05 | 0.74 ± 0.05 | 0.86 |
| Xylazine | 88.9 ± 4.1 | 6.09 ± 0.07 | 0.76 ± 0.05 | 1.60 |
| 3-OH Xylazine | 98.4 ± 1.8 | 6.35 ± 0.03 | 0.98 ± 0.06 | 1.79 |
| 4-OH Xylazine | 83.2 ± 5.6 | 5.64 ± 0.10 | 0.83 ± 0.10 | 2.42 |
| Alpha2B |  |  |  |  |
| Norepinephrine | 107.4 ± 3.7 | 7.90 ± 0.10 | 0.57 ± 0.06 | 0.00 |
| Apraclonidine | 84.3 ± 3.7 | 7.13 ± 0.07 | 0.85 ± 0.09 | 0.66 |
| Clonidine | 82.3 ± 2.6 | 7.04 ± 0.06 | 0.90 ± 0.09 | 0.74 |
| Detomidine | 128.3 ± 1.7 | 8.63 ± 0.04 | 0.78 ± 0.05 | -0.65 |
| Dexmedetomidine | 102.9 ± 0.6 | 9.52 ± 0.02 | 0.83 ± 0.03 | -1.64 |
| Guanfacine | 97.3 ± 7.6 | 6.79 ± 0.17 | 0.63 ± 0.10 | 1.07 |
| Lofexidine | 114.0 ± 2.0 | 8.00 ± 0.05 | 0.78 ± 0.06 | -0.07 |
| Medetomidine | 133.2 ± 1.7 | 8.81 ± 0.04 | 0.98 ± 0.07 | -0.82 |
| Romifidine | 98.7 ± 2.7 | 6.92 ± 0.05 | 0.84 ± 0.07 | 0.94 |
| Tizanidine | 109.3 ± 22.7 | 5.68 ± 0.37 | 0.55 ± 0.09 | 2.23 |
| Xylazine | 88.9 ± 4.1 | 6.10 ± 0.07 | 0.76 ± 0.05 | 1.72 |
| 3-OH Xylazine | 93.3 ± 3.6 | 5.84 ± 0.06 | 0.83 ± 0.07 | 2.00 |
| 4-OH Xylazine | 100.0 ±16.8 | 5.09 ± 0.25 | 0.69 ± 0.10 | 2.78 |
| Alpha2C |  |  |  |  |
| Norepinephrine | 102.0 ± 1.4 | 8.32 ± 0.04 | 0.63 ± 0.03 | 0.00 |
| Apraclonidine | 93.2 ± 1.6 | 7.67 ± 0.04 | 0.69 ± 0.04 | 0.61 |
| Clonidine | 81.9 ± 2.0 | 7.22 ± 0.05 | 0.68 ± 0.04 | 1.00 |
| Detomidine | 104.3 ± 0.9 | 8.94 ± 0.03 | 0.69 ± 0.03 | -0.61 |
| Dexmedetomidine | 107.0 ± 0.6 | 9.75 ± 0.02 | 0.99 ± 0.04 | -1.41 |
| Guanfacine | 91.5 ± 4.9 | 7.03 ± 0.12 | 0.66 ± 0.08 | 1.24 |
| Lofexidine | 99.5 ± 0.9 | 8.56 ± 0.03 | 0.72 ± 0.03 | -0.25 |
| Medetomidine | 105.3 ± 0.9 | 9.25 ± 0.03 | 0.78 ± 0.04 | -0.92 |
| Romifidine | 88.0 ± 2.0 | 7.11 ± 0.05 | 0.76 ± 0.05 | 1.15 |
| Tizanidine | 83.1 ± 4.1 | 6.27 ± 0.08 | 0.78 ± 0.07 | 1.96 |
| Xylazine | 88.2 ± 4.0 | 6.36 ± 0.07 | 0.89 ± 0.09 | 1.87 |
| 3-OH Xylazine | 101.7 ± 1.4 | 7.10 ± 0.03 | 0.76 ± 0.03 | 1.22 |
| 4-OH Xylazine | 106.7 ± 9.1 | 5.38 ± 0.16 | 0.55 ± 0.05 | 2.96 |

While our GPCR Tango study revealed all compounds are active in the ß-arrestin recruitment assays, we note that since all our GPCR Tango constructs have the V_2_-tail fragment to effectively enhance the affinity for ß-arrestin2, we next examined a more selective assay platform. Here we exploited ß-arrestin2 association BRET assays to directly measure the arrestin recruiting activity without the enhanced effect from the V_2_-tail. As shown in Figure 5 and Table 5, ß-arrestin2 association BRET results revealed significant differences of the tested compounds at α_2A_-ARs vs. α_2B_-ARs or α_2C_-ARs. At α_2A_-ARs, apraclonidine, dexmedetomidine, and medetomidine displayed weak partial agonist activity, while other tested agonists had little activity. Detomidine, dexmedetomidine, and medetomidine all were efficacious and potent agonists, while lofexidine had weak partial agonist activity only at α_2B_-ARs and α_2C_-ARsreceptors. Apraclonidine is the only drug with similar activity at all three subtypes. The activity of three drugs (romifidine, tizanidine, and xylazine) for ß-Arrestin2 recruitment was with low efficacy and or potency.

**Fig 5.**
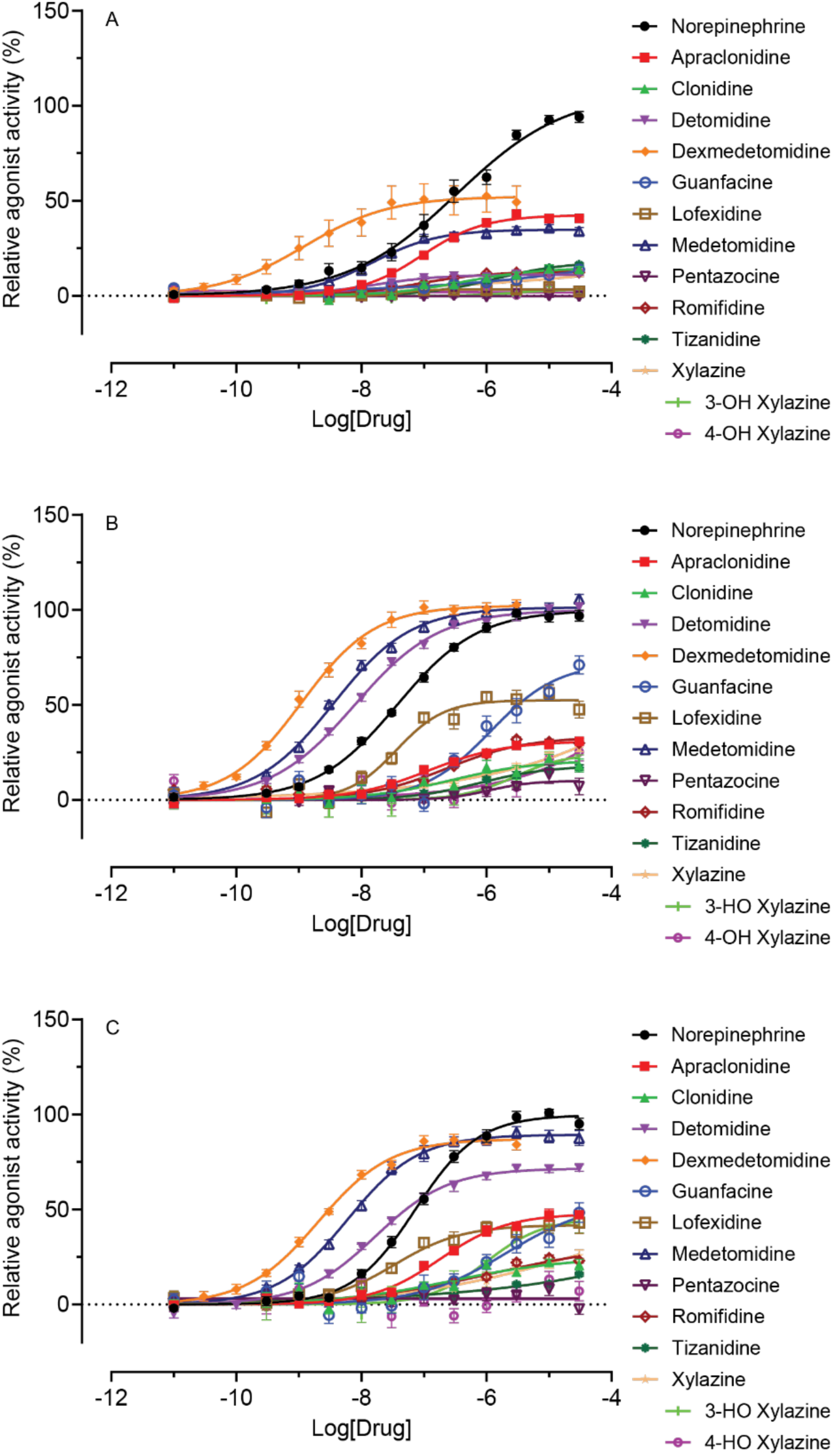
Comparison of agonist activity of FDA approved Alpha2A agonists at Alpha2A (A), Alpha2B(B), and Alpha2C (C) in ß-Arriestin2 association assay (BRET1). The assays were performed in transiently transfected HEK293 T cells. Results were normalized to reference agonist Norepinephrine, represented means ± SEM from a minimum of 3 independent assays, each in quadruplicate set, and pooled for nonlinear fitting to the built-in four-parameter logistic function in Prism (V10). Pharmacological parameters were extracted and summarized in Table 5.

**Table 5.** Agonist activity for ß-Arrestin2 association. Parameters were extracted from nonlinear fitting of concentration response curves as in Figure 5. Note: * E_max_ at 30 μM due to unstable fitting.

| Compounds | E<sub>max</sub> (%) | pEC<sub>50</sub> | Hill | ΔLog(E<sub>max</sub>/EC<sub>50</sub>) |
|---|---|---|---|---|
| Alpha2A |  |  |  |  |
| Norepinephrine | 107.0 ± 6.4 | 6.48 ± 0.15 | 0.50 ± 0.05 | 0.00 |
| Apraclonidine | 42.4 ± 0.8 | 7.04 ± 0.04 | 0.87 ± 0.06 | -0.96 |
| Clonidine | 14.7 ± 1.7 | 6.37 ± 0.24 | 0.72 ± 0.20 | -0.75 |
| Detomidine | 11.1 ± 0.4 | 7.84 ± 0.11 | 0.78 ± 0.13 | -2.34 |
| Dexmedetomidine | 52.2 ± 4.0 | 8.91 ± 0.23 | 0.64 ± 0.18 | -2.74 |
| Guanfacine | 13.8 ± 2.1* | NA | NA | NA |
| Lofexidine | 2.2 ± 1.3* | NA | NA | NA |
| Medetomidine | 34.8 ± 0.8 | 7.83 ± 0.06 | 0.82 ± 0.08 | -1.94 |
| Pentazocine | -0.3 ± 0.7* | NA | NA | NA |
| Romifidine | 12.8 ± 0.6 | 7.09 ± 0.11 | 0.77 ± 0.12 | -1.53 |
| Tizanidine | 17.8 ± 1.7 | 5.86 ± 0.16 | 0.78 ± 0.14 | -0.16 |
| Xylazine | 11.8 ± 1.7* | NA | NA | NA |
| 3-OH Xylazine | 2.4 ± 1.9* | NA | NA | NA |
| 4-OH Xylazine | 1.1 ± 1.04* | NA | NA | NA |
| Alpha2B |  |  |  |  |
| Norepinephrine | 99.8 ± 1.5 | 7.43 ± 0.04 | 0.67 ± 0.03 | 0.00 |
| Apraclonidine | 30.9 ± 1.5 | 6.93 ± 0.12 | 0.71 ± 0.11 | -0.01 |
| Clonidine | 20.8 ± 2.9 | 6.56 ± 0.36 | 0.64 ± 0.24 | 0.19 |
| Detomidine | 99.9 ± 1.2 | 8.11 ± 0.04 | 0.65 ± 0.03 | -0.68 |
| Dexmedetomidine | 102.2 ± 1.6 | 8.96 ± 0.05 | 0.74 ± 0.05 | -1.52 |
| Guanfacine | 72.3 ± 7.6 | 5.93 ± 0.18 | 0.80 ± 0.18 | 1.36 |
| Lofexidine | 52.8 ± 2.0 | 7.44 ± 0.09 | 1.13 ± 0.22 | -0.29 |
| Medetomidine | 101.3 ± 1.3 | 8.45 ± 0.04 | 0.70 ± 0.04 | -1.01 |
| Pentazocine | 10.1 ± 2.4 | 5.96 ± 0.35 | 1.26 ± 1.03 | 0.48 |
| Romifidine | 33.2 ± 1.5 | 6.64 ± 0.10 | 0.68 ± 0.08 | 0.31 |
| Tizanidine | 17.9 ± 2.0 | 6.11 ± 0.20 | 0.79 ± 0.20 | 0.57 |
| Xylazine | 27.1 ± 1.8 * | NA | NA | 2.34 |
| 3-OH Xylazine | 24.6 ± 6.5 | 5.55 ± 0.35 | 1.02 ± 0.56 | 1.27 |
| 4-OH Xylazine | 25.3 ± 9.5* | NA | NA | NA |
| Alpha2C |  |  |  |  |
| Norepinephrine | 99.7 ± 1.7 | 7.14 ± 0.04 | 0.86 ± 0.06 | 0.00 |
| Apraclonidine | 47.7 ± 0.08 | 6.74 ± 0.08 | 0.80 ± 0.09 | 0.08 |
| Clonidine | 24.9 ± 5.0 | 6.56 ± 0.52 | 0.47 ± 0.15 | -0.02 |
| Detomidine | 71.6 ± 1.1 | 7.78 ± 0.04 | 0.74 ± 0.05 | -0.78 |
| Dexmedetomidine | 87.1 ± 1.4 | 8.69 ± 0.05 | 0.75 ± 0.05 | -1.61 |
| Guanfacine | 54.6 ± 13.8 | 5.67 ± 0.47 | 0.62 ± 0.21 | 1.21 |
| Lofexidine | 41.8 ± 2.2 | 7.47 ± 0.14 | 0.80 ± 0.17 | -0.71 |
| Medetomidine | 89.3 ± 1.5 | 8.22 ± 0.05 | 0.77 ± 0.06 | -1.13 |
| Pentazocine | -2.4 ± 2.8* | NA | NA | NA |
| Romifidine | 33.5 ± 6.4 | 5.84 ± 0.52 | 0.38 ± 0.07 | 0.83 |
| Tizanidine | 15.9 ± 1.1 * | NA | NA | NA |
| Xylazine | 25.7 ± 2.9 * | NA | NA | NA |
| 3-OH Xylazine | 45.4 ± 6.3 | 5.99 ± 0.22 | 0.95 ± 0.34 | 2.03 |
| 4-OH Xylazine | 7.0 0 ± 10.4* | NA | NA | NA |

**Table 6.** Transduction coefficients of alpha2 agonists from 4 functional readouts. Parameters were extracted from non-linear fitting to the operational model (PMID **22860188**) with the curves from Figures 1S, 4A, 4B, and 5. Note: ND for not determined.

| Compound | Tango | Gi cAMP | GoA BRET | ß-Arrestin2 BRET |
|---|---|---|---|---|
| Alpha2A receptors |  |  |  |  |
| Norepinephrine | 6.92 ± 0.06 | 8.94 ±0.06 | 8.09 ± 0.03 | 6.54 ± 0.04 |
| Apraclonidine | 8.64 ± 0.03 | 8.09 ± 0.08 | 8.26 ± 0.04 | 5.95 ± 0.10 |
| Clonidine | 7.80 ± 0.06 | 8.77 ± 0.06 | 7.91 ± 0.04 | <4.52 |
| Detomidine | 8.17 ± 0.05 | 8.37± 0.08 | 8.60 ± 0.04 | 5.57 ± 0.31 |
| Dexmedetomidine | 6.56 ± 0.18 | 10.13 ± 0.09 | 9.45 ± 0.03 | 8.03 ± 0.08 |
| Guanfacine | 7.17 ± 0.08 | 8.66 ± 0.08 | 7.95 ± 0.04 | <4.52 |
| Lofexidine | 8.25 ± 0.05 | 8.23 ± 0.08 | 8.46 ± 0.04 | <4.52 |
| Medetomidine | 8.75 ± 0.06 | 9.10 ± 0.07 | 9.11 ± 0.03 | 6.57 ± 0.11 |
| Romifidine | 7.75 ± 0.05 | 7.62 ± 0.09 | 7.84 ± 0.04 | 4.94 ± 0.28 |
| Tizanidine | 6.92 ± 0.06 | 6.99 ± 0.09 | 7.31 ± 0.04 | <4.52 |
| Xylazine | 5.04 ± 0.15 | 6.80 ± 0.09 | 6.51 ± 0.04 | 6.44 ± 0.64 |
| 3-OH Xylazine | ND | 6.47 ± 0.07 | 6.23 ± 0.04 | <4.52 |
| 4-OH Xylazine | ND | 5.83 ± 0.07 | 5.42 ± 0.05 | <4.52 |
| Alpha2B receptors |  |  |  |  |
| Norepinephrine | 7.04 ± 0.06 | 8.98 ± 0.05 | 7.82 ± 0.04 | 7.38 ± 0.04 |
| Apraclonidine | 7.81 ± 0.05 | 7.42 ± 0.07 | 6.79 ± 0.06 | 5.95 ± 0.11 |
| Clonidine | 6.98 ± 0.08 | 7.39 ± 0.06 | 6.67 ± 0.06 | 5.34 ± 0.19 |
| Detomidine | 8.45 ± 0.05 | 9.20 ± 0.06 | 8.58 ± 0.03 | 8.06 ± 0.03 |
| Dexmedetomidine | 6.60 ± 0.14 | 9.86 ± 0.05 | 9.30 ± 0.04 | 8.93 ± 0.04 |
| Guanfacine | 5.89 ± 0.12 | 7.02 ± 0.06 | 6.62 ± 0.05 | 5.55 ± 0.06 |
| Lofexidine | 8.14 ± 0.05 | 8.28 ± 0.06 | 7.86 ± 0.04 | 6.79 ± 0.07 |
| Medetomidine | 8.79 ± 0.07 | 9.58 ± 0.06 | 8.79 ± 0.03 | 8.41 ± 0.03 |
| Romifidine | 7.08 ± 0.07 | 6.82 ± 0.08 | 6.68 ± 0.05 | 5.70 ± 0.11 |
| Tizanidine | 6.05 ± 0.09 | 7.41 ± 0.10 | 5.79 ± 0.07 | 4.75 ± 0.18 |
| Xylazine | 4.50 ± 0.23 | 5.90 ± 0.07 | 5.62 ± 0.05 | 4.96 ± 0.15 |
| 3-OH Xylazine | ND | 5.76 ± 0.06 | 5.57 ± 0.06 | <4.52 |
| 4-OH Xylazine | ND | 5.18 ± 0.07 | 4.94 ± 0.07 | <4.52 |
| Alpha2C receptors |  |  |  |  |
| Norepinephrine | 6.77 ± 0.05 | 8.54 ± 0.06 | 8.26 ± 0.03 | 7.12 ± 0.04 |
| Apraclonidine | 6.91 ± 0.06 | 6.69 ± 0.10 | 7.51 ± 0.03 | 6.17 ± 0.08 |
| Clonidine | 5.61 ± 0.13 | 6.37 ± 0.10 | 6.97 ə 0.04 | 5.73 ± 0.19 |
| Detomidine | 7.46 ± 0.06 | 7.80 ± 0.10 | 8.88 ± 0.03 | 7.49 ± 0.05 |
| Dexmedetomidine | 6.84 ± 0.14 | 9.01 ± 0.07 | 9.70 ± 0.02 | 8.55 ± 0.05 |
| Guanfacine | 5.85 ± 0.07 | 6.49 ± 0.07 | 6.87 ± 0.04 | 5.35 ± 0.10 |
| Lofexidine | 7.14 ± 0.08 | 7.75 ± 0.08 | 8.45 ± 0.03 | 6.81 ± 0.10 |
| Medetomidine | 7.88 ± 0.06 | 8.55 ± 0.08 | 9.18 ± 0.03 | 8.10 ± 0.04 |
| Romifidine | 6.09 ± 0.09 | 5.64 ± 0.12 | 6.89 ± 0.04 | 5.49 ± 0.15 |
| Tizanidine | 4.70 ± 0.15 | 5.42 ± 0.16 | 6.00 ± 0.05 | 4.89 ± 0.24 |
| Xylazine | 4.48 ± 0.16 | 5.33 ± 0.09 | 6.18 ± 0.04 | 5.24 ± 0.18 |
| 3-OH Xylazine | ND | 6.31 ± 0.07 | 7.01 ± 0.03 | 5.37 ± 0.12 |
| 4-OH Xylazine | ND | 4.89 ± 0.11 | 5.50 ± 0.04 | 7.60 ± 1.90 |

## Discussion

The main finding of this paper is that xylazine differs from other tested and FDA approved α_2_-AR agonists by being a moderately potent k-opioid receptor (KOR) agonist and a highly G protein biased α_2_-AR agonist. Other tested α_2_-AR agonist displayed variable on- and off-target activities at a variety of tested GPCRs and these activities could be involved in their actions, although further studies will be needed to determine if these activities are clinically relevant.

Of interest, we found that the tested α_2_-AR agonists had differential activities in β-arrestin-recruitment assays. Most of the tested α_2_-AR agonists were low-efficacy partial agonists at the α_2A_-AR and α_2C_-AR, both of which regulate sympathetic neurotransmission ^15^ Our data demonstrates that dexmedetomidine is the most efficacious of the tested α_2_-AR agonists for β-arrestin recruitment, and xylazine is the least. This differential ability of tested α_2_-AR agonists for β-arrestin recruitment is potentially important two reasons: Firstly, Dexmedetomidine is being used to manage xylazine withdrawal in clinical settings^16^; and secondly Dexmedetomidine/medetomidine is now increasingly observed in the unregulated drug supply itself ^17^ requiring additional understanding of how these drugs signal to consider treatment options. As the unregulated drug supply transitions from xylazine to (dex)medetomidine the differences in signaling and potency of these compounds, how they impact arrestin recruitment *in vivo* and alter tolerance, and how they potentially synergize with opioids will need to be closely examined across systems.

Currently, it is unknown what the actual concentrations of xylazine are among individual who are abusing xylazine/fentanyl mixtures although post-mortem blood samples provide concentrations from the periphery ranging from 5.2-200 μg/l (.02-0.9 μM) ^18^ and 3.3-2,755 μg/l (0.015 – 13.8 μM) ^19^. As xylazine’s agonist potency at KOR against GoA (the endogenous G protein) is 2.3 μM, actions of xylazine at KOR are within the reported post-mortem concentration ranges. With regard to these studies, we note that there is scant information as to when these samples were retrieved relative to death. The methods state simply that the results were analyzed ‘within 72 hours ‘of sample submission and the median time from date of death to results reporting to the medical examiner office was 7 days. A clinical study from Lin et al. ^20^in living patients observed blood xylazine concentrations as high as

127.2 µg/L (0.6 μM) however it was unknown when the last drug administration was prior to blood sampling. Extrapolating from the these studies it is clear that xylazine may achieve concentrations sufficient to activate κOR receptors. Data from male rats has demonstrated that a single IV bolus injection of xylazine rapidly (within 15 minutes) concentrates in the brain vs. the blood and is an order of magnitude higher in the brain vs. plasma for upwards of 4 hours, ^21^this study supports findings from others that demonstrate that due xylazine’s lipophilic nature it reaches highest concentrations in the brain and kidneys ^22^. These data suggest that blood concentrations may not accurately reflect the quantity and molarity of xylazine that the brain is exposed to, and that researchers should consider collecting brain samples.

Our finding that xylazine has activity at κORs, like pentazocine, is intriguing because both drugs are misused by people and are well known to be associated with skin lesions^5,6^. Pentazocine, often combined with the antihistamine tripelennamine are commonly referred to as the “T’s and Blues”. Pentazocine misuse was first reported in the medical community before becoming a common street drug in the 1980s^23^. Therefore, their similar action at the KOR may represent an avenue by which both drugs are potentially causing necrotic skin lesions, as human skin cells are known to express KOR^24^. Indeed, a recent study has suggested that xylazine associated skin lesions are more likely due to “skin popping” or subcutaneous injection^6^.

As we have previously demonstrated, many CNS-active medications including antipsychotics^25^, antidepressants ^25^ ^26^, psychedelics ^27^, anti-Parkinsonian medications ^28^, entactogens ^29^, anorectics^30^ and anxiolytics ^31^ have potent off-target actions. These off-target actions have been implicated in antipsychotic drug-induced weight gain ^32^ and drug-induced valvular heart disease, for example ^33^ Given the differential on- and off-target *in vitro* pharmacology of the tested α_2_-AR agonists, it is conceivable that these activities could differentially affect brain circuits involved in attention, arousal, affective behavior, reward, and stress. In future studies it will be important to determine if such signaling differences are also observed in native expression environments and if they are behaviorally relevant. Taken together, our findings indicate that α_2_-AR agonists, like many other CNS active medications, display complex on- and off-target pharmacological profiles and that future studies should investigate the potential relevance of these activities for both their therapeutic actions and side-effects.

## Materials and Methods

### Materials

Compounds were purchased from MCE, Cayman, or Tocris. Human GPCR constructs were reported before (PMID **25895059**) or subcloned from corresponding Tango constructs, and all available from AddGene.

### PRESTO-Tango GPCRome assays

GPCRome screening assays were carried out according to previously published procedure (PMID **25895059**) with modifications as detailed below. In detail, HTLA cells were maintained in regular growth medium (DMEM, 10% FBS, and Pen/Strep (100 Units/ml penicillin, and100 ug/ml streptomycin)) supplemented with 100 ug/ml Hygromycin B and 15 ug/ml Blasticidin. For screening, the HTLA cells were seeded in PLL coated 384-well white clear-bottom plates in the DMEM with 1% dialyzed FBS (dFBS) and Pen/Strep at the density of 10,000 cells per well per 40 µl. After recovery for about 5 hours in the cell incubator, the cells were transfected using a modified PEI transfection protocol. Each sample needs 8x 384-well cell plates and we usually run the screening in a batch of 10 to 15 samples. For a set of 10 samples, DNA constructs were prepared in 96-well drug plates (2.5 µg in 5 µl/well) and diluted with 180 µl Opti-MEM, followed by 12.5 µl PEI transfection reagent. The DNA plate received another 180 µl Opti-MEM to bring the final volume to 365 µl per well. The second addition of 180 µl Opti-MEM served as a mix purpose. The DNA plate was then incubated at the room temperature for 30 min. The mixture was then aliquoted and transferred into 384-well drug plates at 40 µl/well to mix with pre-added 60 µl/well Opti-MEM. Transfection was performed on the Hamilton Star robot system with a 384-well head by transferring 6 µl/well of DNA mix into HTLA cells plates. The final DNA amount is about 16 ng per well and each DNA construct was transfected in a total of 8 wells. Dopamine D2 receptor served as an assay control and was transfected at about 8 ng per well for a total of 2 columns of 32 wells. Overnight incubation after transfections, samples were added to the cells at 10 µl per well by the Hamilton Star robot. Each construct (8 wells) has 4 wells for samples and 4 wells for basal with medium alone. The cells were stimulated overnight. To count luminescence activity in the SpectraMax L (Molecular Devices), media and drug solutions were removed and 20 µl/well 20x diluted BrightGlo reagent (Promega) with drug buffer (1x HBSS, 20 mM HEPES, pH 7.4) was added for counting after 20 min incubation at room temperature in the dark. Results were calculated in the form of fold of average basal for each construct.

### GPCR Tango concentration response assays

The HTLA cells were maintained as above. One day before transfection with individual DNA construct, the HTLA cells were plated in the regular growth medium without Hygromycin or Blasticidin (such as 1.5 million cells per 6-cm dish) and then transfected using a modified PEI procedure. Briefly, DNA (2 – 3 µg) was diluted into 150 µl Opti-MEM, followed by addition of 10-15 ul PEI solution. The mixture was incubated for about 30 min at room temperature before adding to the HTLA cells. The overnight transfected cells were plated into PLL-coated white clear bottom 384-well plates at the density of 10,000 cells per well per plate at the density of 10,000 cells per well per 40 µl in DMEM with 1% dFBS and Pen/Strep. Drugs were added after plated cells were recovered for about 5 hours. Sample dilutions were made in the same DMEM with 1% dFBS and Pen/Strep at 5x and added to cells at 10 µl per well. Stimulation was kept overnight before counting as above. Results normalized and analyzed in the GraphPad Prism V10.

### GloSensor cAMP assays

GloSensor cAMP assays were conducted in transiently transfected HEK293 T cells. Briefly, HEK293 T cells were maintained and transfected in regular growth medium with receptor DNA and GloSensor cAMP plasmid (Promega) as above using the PEI protocol. Overnight transfected cells were plated in PLL coated 384-well white clear bottom plates using DMEM with 1% dFBS and Pen/Strep at 10,000 cells per well per 40 µl. After at least 5 hours (up to overnight) recovery, medium was removed and the cells received 20 µl/well drug solution at 1.5x of final concentrations. The drug solutions were prepared in the assay buffer (1x HBSS, 20 mM HEPES, pH 7.4, 0.5 mg/ml BSA) supplemented with 35 mM Luciferin. After 20 min incubation at room temperature in the dark, Luminescence was read in the SpectraMax L (Molecular Devices) for stimulatory activity (G_αs_ mediated cAMP production). Right after the first reading, the plate received 10 µl Isoproterenol (ISO) at 300 nM (100 nM final) to activate endogenous ß_2_ receptors to increase adenylyl cyclase activity. The G_i/o_ activity was read 20 min after ISO addition and incubation. Control assays were performed side by side using HEK293 T cells transfected with GloSensor cAMP plasmid alone to determine potential nonspecific activity. Results were normalized and analyzed in the GraphPad Prism V10.

### Bioluminescence Resonance Energy Transfer (BRET) assays

BRET assays were conducted according to previously published protocol (**32367019 and 36917754**) with modifications. In detail, HEK293 T cells were maintained and transfected in regular growth medium using the PEI method as above. For a 10-cm dish, the total DNA amount is 8 µg. For ß-Arrestin2 association BRET1 assays, HEK293 T cells were transfected with GPCR-rLuc and ß-Arrestin2-Venus constructs at the ratio of 1:2. For G protein dissociation BRET2 assays, HEK293 T cells were transfected with receptor and G protein subunits (G_α_-rLuc, G_ß_, and G_Γ_-GFP) at ratio of 1:1:1:1. Overnight transfected cells were plated in PLL-coated 384-well plates using DMEM with 1% dFBS and Pen/Strep at the density of 10,000 cells per well per 40 µl. After at least 5 hours (up to overnight) recovery, medium was removed and the cells received 20 µl/well drug solutions at 1.5x of final concentrations. The drug solutions were prepared in the assay buffer (1x HBSS, 20 mM HEPES, pH 7.4, 0.5 mg/ml BSA). After incubation at room temperature and in the dark for 10 minutes, 10 µl/well luciferase substrate was added at the final of 5 µM (Coelenterazine h for ß-Arrestin2 association BRET1 assays and Coelenterazine 400a for G-protein dissociation BRET2 assays) at timed steps (1.5 minutes for every 4 rows). The plate was read in the PHERAStar FSX (BMG LABTECH) at 5, 10, and 15 minutes after substrate addition. Readings at 5 or 10 minutes are usually used for data analysis in Prism V10.

## ACKNOWLEDGEMENTS

XPH and BLR are supported by the National Institute of Mental Health Psychoactive Drug Screen Program (NIMH PDSP), contract # 75N95023C00021. The NIMH PDSP is directed by Bryan l Roth, MD, PhD, at the University of North Carolina at Chapel Hill and Project Officer Jamie Driscoll at NIMH, Bethesda, MD. BK and BLR were supported by RO1MH122205.

This project is also supported by the Food and Drug Administration (FDA) of the U.S. Department of Health and Human Services (HHS) as part of a financial assistance award for Triangle Center of Excellence in Regulatory Science & Innovation, U01FD007857, totaling $1,453,116 from Center for Drug Evaluation and Research (CDER). The contents are those of the author(s) and do not necessarily represent the official views of, nor an endorsement, by FDA/HHS, or the U.S. Government.

